# *In vivo* Antidiarrheal activities of the hydro alcoholic extracts of *Schinus molle* L. (Anarcardiaceae) leaf in mice

**DOI:** 10.1101/2022.07.06.499014

**Authors:** Getnet Mengistu, Kidan Hailay, Desye Misganaw, Abel Andualem, Yaschilal Muche Belayneh

## Abstract

**Background:** diarrhea is the second leading cause of death among children under five globally next to pneumonia and it kills around 760, 000 children every year in the world. The available synthetic drugs to treat diarrhea have limited safety and efficacy. Thus, people are still dependent on the herbal remedies for managing diarrhea.

**Objective:** To evaluate in vivo antidiarrheal activities of the hydro-alcoholic leaf extracts of *Schinus molle* L. (Anacardiaceae) in mice.

**Methods:** The 80% methanol leaf extract was prepared by successive maceration technique. The antidiarrheal activity of the extract was evaluated using castor oil induced diarrhea, enteropooling and small intestine transit models. The test groups received various doses (100, 200 and 400 mg/kg) of the extract, whereas positive controls received Loperamide (3 mg/kg) and negative controls received vehicle (distilled water, 10 ml/kg). Data were analysed using one-way ANOVA followed by Tukey’s posthoc test.

**Result:** In the castor oil induced diarrheal model, the 80% methanol extract delayed onset of defecation and reduced the number and weight of feces at all tested doses significantly as compared to the negative control. In the enteropooling test, the 80ME significantly (P<0.001) reduced the weight and volume of intestinal fluid at all tested doses when compared to negative control. Results from the charcoal meal test revealed that the extracts produced a significant anti-motility effect at all tested doses as compared to negative control.

**Conclusion:** This study confirmed the antidiarrheal activity of the hydro-alcoholic extract and the highest test dose produced the maximum antidiarrheal activity in all models

## Introduction

Diarrhea is the passage of three or more loose or liquid stools per day (or more frequent passage than is normal for the individual) due to an alteration in a normal bowel movement (1, 2). Since it is difficult to quantitate consistency of the stool and the weight is affected by the type of food consumed, a combination of frequency, stool consistency and stool weight should be taken into account for assessing diarrhea (3).

Based on duration of symptoms, diarrhea can be classified into acute, persistent and chronic diarrhea. Diarrhea lasting less than 2 weeks is considered acute which is typically self-limiting and resolves quickly with no lasting sequelae. Persistent diarrhea starts suddenly as acute diarrhea with duration varying from 2 to 4 weeks and chronic diarrhea lasts longer than four weeks (4).

Even though diarrhea is a preventable and treatable disorder, it remains the 2^nd^ leading cause of mortality among children under five years of age and responsible for killing around 760,000 children every year (5, 6). Of all child deaths from diarrhea, 78% occur in the African and South-East Asian regions (5, 7) and Ethiopia’s diarrhea mortality rate is the 5^th^ highest in the world and diarrhea is the 2^nd^ leading cause of death across all ages in Ethiopia (7, 8).

Most of the drugs used for the management of diarrhea are not effective and associated with different side effects. Moreover, there is an increasing threat of drug resistance, side effects, superinfection and the possibility of induction of disease producing bacteriophages by antibiotics. Due to these problems, WHO encourages studies for the treatment and prevention of diarrheal diseases depending on traditional medical practices. Therefore, treating diarrhea with plant-derived compounds, which are accessible and do not require sophisticated pharmaceutical synthesis seems highly attractive (9, 10).

Schinus molle L. is traditionally used for the treatment of hepatic disease/jaundice and diarrhea in Ethiopia. An ethnobotanical survey studies done in Ethiopia reported that the leaf juice of Schinus molle is taken orally to treat diarrhea (11, 12). However; the antidiarrheal activity of this plant has not been scientifically studied. Additionally, different previous experimental studies have reported that *Schinus molle* have significant *in vitro* antioxidant, antibacterial and anti-proliferative and *in vivo* anti-inflammatory activities (13–16) and phytochemical studies showed the plant is rich in phenols, flavonoids, tannins and anthocyanins (14, 16), suggesting the plant may have antidiarrheal effect. Therefore, intensive research has to be conducted on the traditionally used medicinal plants for the purpose of developing new, effective and safe antidiarrheal drugs. So, this study was undertaken to evaluate in vivo antidiarrheal activities of the hydro-alcoholic extracts of *Schinus molle* L. (Anacardiaceae) leaf in mice.

## Materials and Methods

### Chemicals and reagents

The main chemicals that were used include distilled water, 2% Tween80, 80% methanol, n-butanol, chloroform, CCl_4_, 10% formalin, ether, normal saline, paraffin wax, hematoxylin, eosin, xylol, 2,2-diphenyl-1-picrylhydrazyl (DPPH), the standard drug Silymarin (Silybon-140), castor oil, activated charcoal, loperamide and reagents used for phytochemical screening. Analytical grade chemicals were used.

### Plant material

The leaves of *Schinus molle* were collected from its natural habitat around Kobbo, North Wollo, North East Ethiopia. Identification and authentication of the plant specimen was done by a taxonomist at the National Herbarium, Department of Biology, Addis Ababa University, where a voucher specimen was deposited for future reference.

### Experimental animals

Healthy adult male and female Swiss albino mice with the age of 6-8 weeks old and weighing 22-30g were maintained in the animal house facilities at department of pharmacy, Wollo University. Female mice were used for acute toxicity test and male mice for the main study. The mice were housed in polypropylene cages (6 mice per cage) under standard environmental conditions and 12-12 hour light-dark cycle and were left for a week for the purpose of acclimatization. They were provided a laboratory pellet diet and water *ad libtum*.

### Preparation of plant extracts

The leaves of the plant were thoroughly washed with tap water to remove dirt and cleaned with gauze. Then, it was cut into pieces manually and dried under shade. The leaves were pulverized by using mortar and pestle to get coarse powder. The powder was extracted by cold maceration technique with 80% methanol to get the crude hydro-alcoholic extract for 72 hours at room temperature. Then, 80% methanol liquid extract was evaporated in order to remove the methanol under vacuum at 40 °C by using a rotary evaporator and the extract was frozen in a refrigerator and dried in a lyophilizer. Finally, the extract were transferred into vials and kept in a desiccator until used for the proposed experiment.

### Acute oral toxicity test

Acute oral toxicity test for the extracts of the leaf of *Schinus molle* was performed according to the organization for economic co-operation and development guideline. Mice were fasted for 3-4 h before and 1-2 h after administration of the extract. First, a sighting study was performed to determine the starting dose. Female mice were used and each mouse was given 2000 mg/kg of the extract as a single dose by oral gavage. Since mortality was not observed within 24 h, additional four mice were used and administered at the same dose mentioned above. The animals were observed continuously for 4 h with 30 min interval and then for 14 consecutive days with an interval of 24 h for the general signs and symptoms of toxicity (diarrhea, weight loss, tremor, lethargy and paralysis), food and water intake and mortality. Then, three dose levels were chosen for the extract: a middle dose, which is one-tenth of the limit dose during acute toxicity study; a low dose, which is half of the middle dose (1/20^th^ of the limit dose) and a high dose which is twice the middle dose (1/5^th^ of the limit dose) (17).

### Determination of antidiarrheal activity

#### Animal grouping and dosing

In all models, animals were randomly divided into five groups (negative control, positive control and three test groups) comprising of six animals per group. Negative controls received distilled water (10 ml/kg) and positive controls received Loperamide (3mg/kg) in all models. The test groups (group 3, 4 and 5) were received different doses (100mg, 200mg & 400mg respectively) of the extract orally which was determined based on the acute oral toxicity test and pilot study.

#### Castor oil induced diarrheal model

The method described by Igboeli *et al* (18) was followed for this study. Swiss albino mice of either sex were fasted for 18 h with free access to water and grouped and treated as described under animal grouping and dosing section. One hour after dosing, each mouse was given 0.5 ml of castor oil orally for induction of diarrhea and placed individually in cages in which the floor is lined with white paper. The transparent paper was changed every hour for a total of four hours. During the observation period, the onset of diarrhea, number and weight of wet stools, total number of feces and total weight of fecal output was recorded. Finally, percentage of fecal output and diarrheal inhibition were calculated by using the formulas described below.

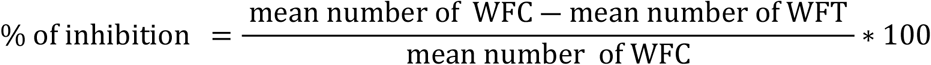

Where, WFC = wet feces in control group and WFT = wet feces in test group

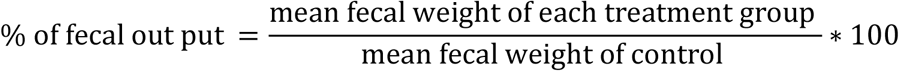

#### Castor oil induced enteropooling test

The effects of the extract on intra-luminal fluid accumulation was determined using the method described by Robert *et a l*(19). Animals were fasted for 18 h, grouped and treated as described under animal grouping and dosing section. After 1 h of treatment, 0.5 ml of castor oil was administered and animals were sacrificed 1 h following castor oil administration. The abdomen of each animal was then opened; the small intestine was ligated at both the pyloric sphincter and the ileo-cecal junction, and dissected. The dissected small intestine was weighed and intestinal contents was then collected by milking into a graduated tube and volume of the contents was measured. Weight of the intestine after milking was taken and the difference between the two weights was then recorded. Finally, percentage of reduction of intestinal secretion (volume and weight) was calculated relative to the negative control using the following formula

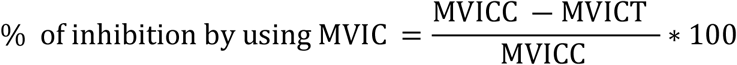

Where, MVIC – Mean Volume of Intestinal Content

MVICC - Mean Volume of Intestinal Content of Control Group

MVICT - Mean Volume of Intestinal Content of Test Group

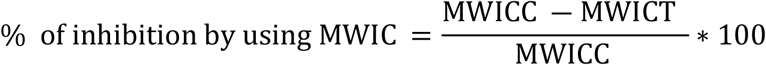

Where, MWIC – Mean Weight of Intestinal Content

MWICC- Mean Weight of Intestinal Content of Control Group

MWICT - Mean Weight of Intestinal Content of Test Group

#### Gastrointestinal motility test

Animals were fasted for 18h with free access to water and divided and treated as described under animal grouping and dosing section one hour before administration of 0.5 ml castor oil. One ml of the marker (5%activated charcoal suspension in water) was administered orally 1 h after castor oil treatment. The animals were sacrificed 1 h after charcoal meal and the small intestine dissected out from pylorus to caecum and placed length wise on a white paper. The distance travelled by the marker and the total length of the intestine was measured. The peristaltic index and percentage of inhibition were calculated by using the following formula (18, 20).

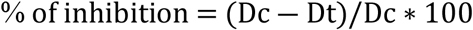

Where, Dc: Mean distance travelled by the charcoal in the control group and

Dt: Mean distance travelled by the charcoal in the test group

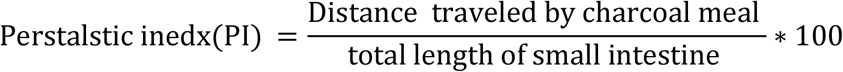

#### In-vivo antidiarrheal index

The *in vivo* antidiarrheal index (ADI) for the positive control and different doses of the extract was determined based on the data from the above tests using the formula developed by Aye-Than *et al* (20).

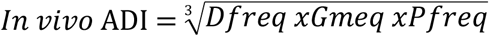

Where, Dfreq is the delay in defecation time as % of negative control, Gmeq is the gut meal travel reduction as % of negative control and Pfreq is the reduction in the number of stools as % of negative control.

#### Phytochemical screening

Phytochemical screening tests were carried out for the extract by using standard procedures to identify the presence of secondary metabolites like tannins, saponins, flavonoids, terpenoids, steroids, alkaloids and cardiac glycosides (21, 22).

#### Statistical analysis

Results of the study were expressed as mean ± standard error of mean (S.E.M). Statistical package for social science (SPSS) version 23 was used to analyse the results. Statistically significant differences between groups were evaluated by analysis of variance (one-way ANOVA) followed by Tukey’s multiple comparison test. The analysis was performed with 95% confidence interval and probabilities less than 0.05 (p<0.05) was considered significant.

## Result

### Acute oral toxicity test

The acute oral toxicity test of 80ME of leaves of Schinus molle L. indicated that it didn’t cause gross behavioural changes and mortality within 24 h as well as in the next 14 days, indicating that the LD50 of the extract was greater than 2000 mg/kg in mice.

### Effects of 80% methanol extract on castor oil induced diarrhea

In the castor oil induced diarrheal model, 80ME of leaves of Schinus molle L. delayed the onset of defecation and reduced the frequency of defecation at all tested doses (100, 200 & 400mg/kg) significantly (P<0.001) as compared to the negative control. Results from the experiment also revealed that all the tested doses of 80ME significantly (p<0.001) reduced the weight of feces (wet feces and total feces) when compared with negative control as depicted in Table 1.

**Table 1:** Effects of 80% methanol leaf extract of Schinus molle L. on castor oil induced diarrhea in mice

| Group | Onset of diarrhea (min) | No of wet faces | Total no of faces | Weight of wet faces | Weight of total faces |
| --- | --- | --- | --- | --- | --- |
| Control | 13.2 ± 1.08 | 6.6 ± 0.33 | 9.7 ± 0.33 | 0.39 ± 0.01 | 0.46 ± 0.01 |
| 80ME100 | 35.7 ± 1.50<sup>a3b3c3d3</sup> | 3.1 ± 0.31<sup>a3b2c2d3</sup> | 6.8 ± 0.31<sup>a3b3c3d3</sup> | 0.24 ± 0.02<sup>a3b3c3d3</sup> | 0.34 ± 0.02<sup>a3b3c3d3</sup> |
| 80ME200 | 68.3 ± 2.10<sup>a3b3d3</sup> | 1.6 ± 0.21<sup>a3</sup> | 4.5 ± 0.22<sup>a3b3d3</sup> | 0.14 ± 0.02<sup>a3d3</sup> | 0.22 ± 0.02<sup>a3b3d3</sup> |
| 80ME400 | 185.3 ± 3.72<sup>a3b3</sup> | .8 ± 0.17<sup>a3</sup> | 2.3 ± 0.21<sup>a3b1</sup> | 0.04 ± 0.01<sup>a3</sup> | 0.08 ± 0.01<sup>a3</sup> |
| Loperamide | 93.3 ± 2.40<sup>a3b3</sup> | 1.5 ± 0.22<sup>a3</sup> | 3.5 ± 0.22<sup>a3</sup> | 0.08 ± 0.01<sup>a3</sup> | 0.13 ± 0.01<sup>a3</sup> |

Data are expressed as mean ± SEM (n=6); analysis was performed with One-Way ANOVA followed by Tukey test; <sup>a</sup> compared to negative control; <sup>b</sup> compared to Loperamide, 3mg/kg; <sup>c</sup> compared to 200 mg/kg; <sup>d</sup> compared to 400 mg/kg; <sup>1</sup>p<0.05, <sup>2</sup>p<0.01, <sup>3</sup>p<0.001; 80ME, 80% methanol extract, Negative controls received distilled water.

The highest tested dose of 80ME (400 mg/kg) showed the maximum percentage inhibition of defecation and the lowest percentage of mean fecal output when compared with the tested doses of the extract and positive control as shown on fig. 1.

**Fig 1.**
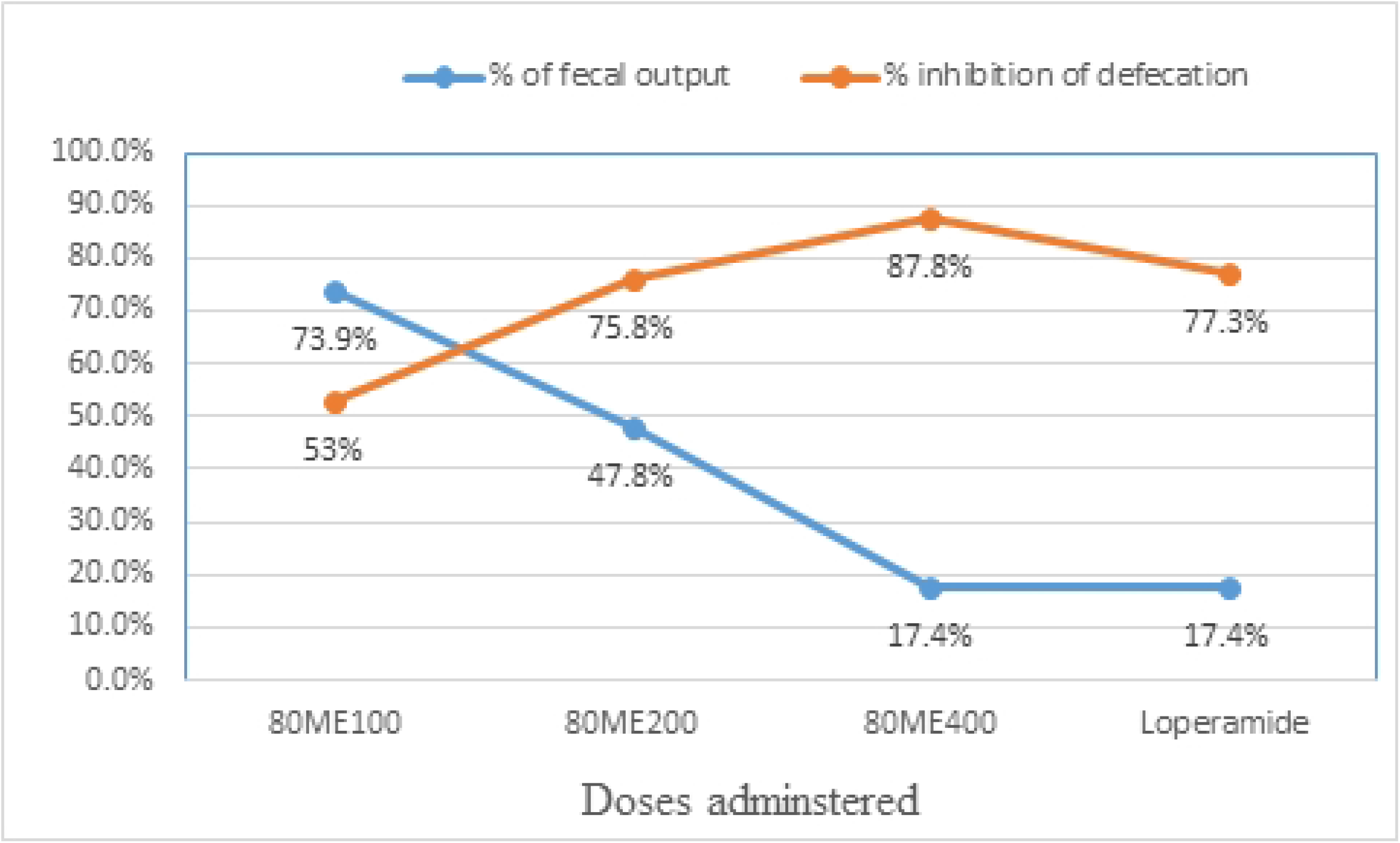
Percentage inhibition of defecation and % of fecal output of the 80% methanol leaf extract of Schinus molle L. on castor oil induced diarrhea in mice

### Effects of 80% methanol extract on castor oil-induced enteropooling

In the gastrointestinal enteropooling test, the 80ME of leaves of Schinus molle L. significantly reduced the weight and volume of intestinal content at all tested doses of the extract (P<0.001) when compared to the negative control. The highest effect on reducing both weight and volume of intestinal content was achieved by the highest doses of the extract (400mg) as shown on table 2.

**Table 2:**
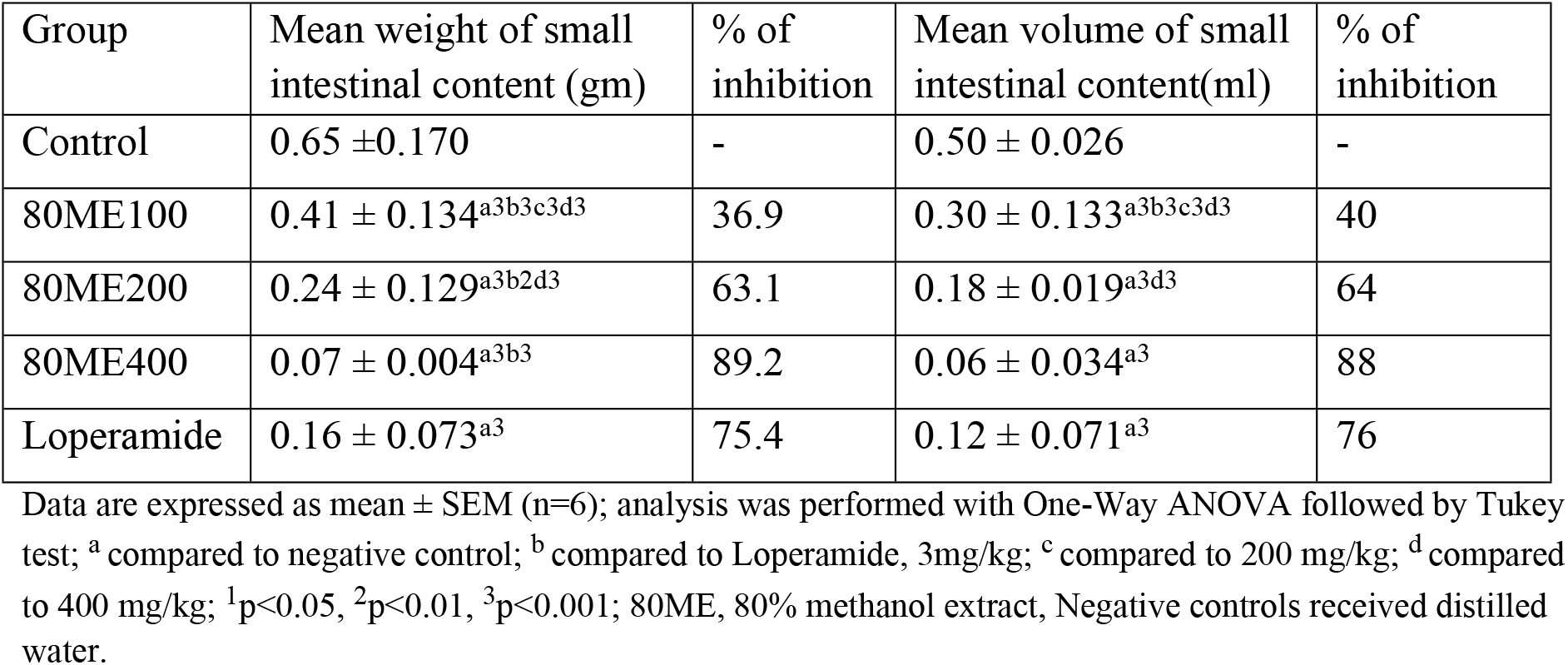
Effects of 80% methanol leaf extract of Schinus molle L. on castor oil induced enteropooling in mice

### Effect of 80% methanol extract on gastrointestinal motility

The 80% methanol extract reduced castor oil induced movement significantly at all doses (p < 0.001) when compared with negative control and the maximum effect is obtained by 400 mg/kg of the tested dose (73.7%) as shown in Tables 3.

**Table 3:**
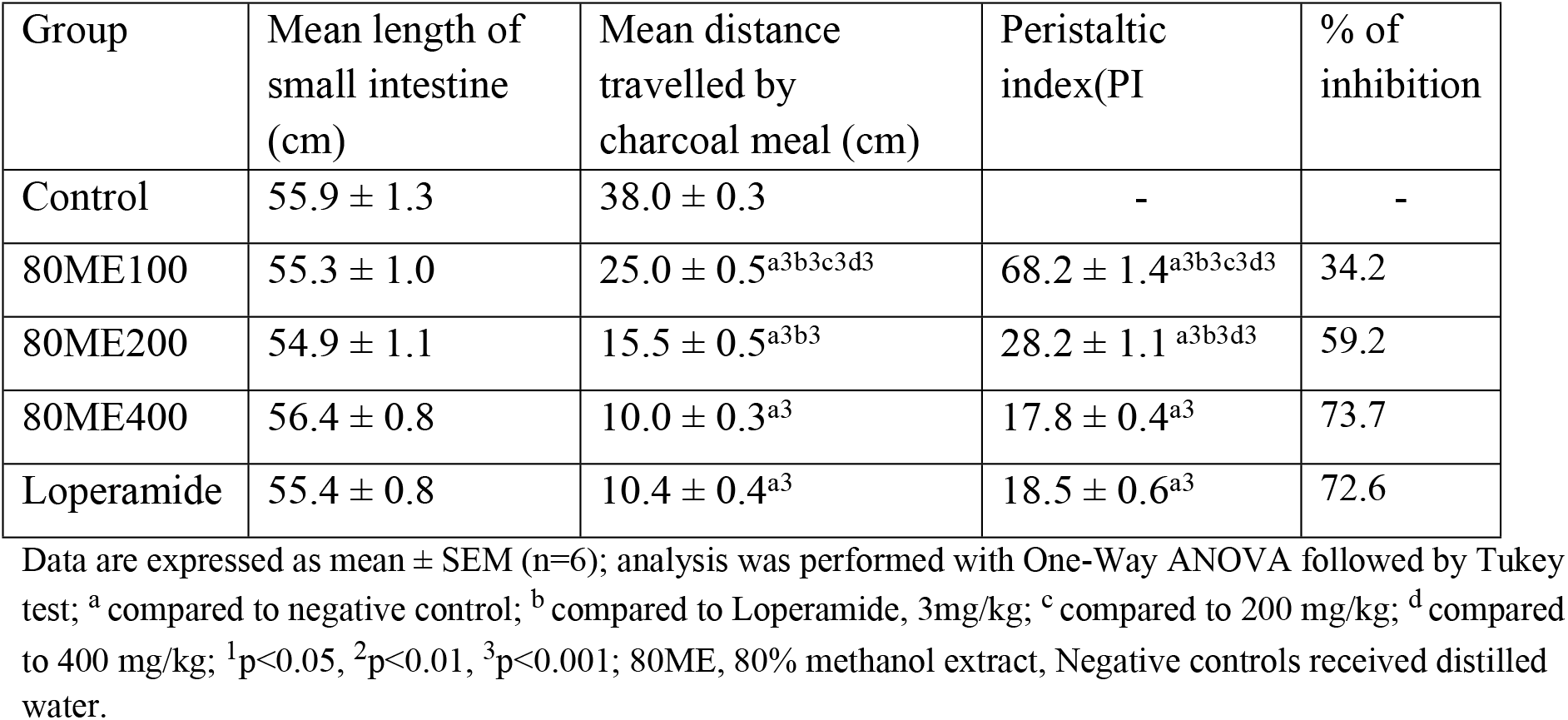
Effects of 80% methanol leaf extract of Schinus molle L. on castor oil induced gastrointestinal motility in mice

| Group | Mean length of small intestine (cm) | Mean distance travelled by charcoal meal (cm) | Peristaltic index(PI | % of inhibition |
| --- | --- | --- | --- | --- |
| Control | 55.9 ± 1.3 | 38.0 ± 0.3 | - | - |
| 80ME100 | 55.3 ± 1.0 | 25.0 ± 0.5<sup>a3b3c3d3</sup> | 68.2 ± 1.4<sup>a3b3c3d3</sup> | 34.2 |
| 80ME200 | 54.9 ± 1.1 | 15.5 ± 0.5<sup>a3b3</sup> | 28.2 ± 1.1<sup>a3b3d3</sup> | 59.2 |
| 80ME400 | 56.4 ± 0.8 | 10.0 ± 0.3<sup>a3</sup> | 17.8 ± 0.4<sup>a3</sup> | 73.7 |
| Loperamide | 55.4 ± 0.8 | 10.4 ± 0.4<sup>a3</sup> | 18.5 ± 0.6<sup>a3</sup> | 72.6 |

Data are expressed as mean ± SEM (n=6); analysis was performed with One-Way ANOVA followed by Tukey test; <sup>a</sup> compared to negative control; <sup>b</sup> compared to Loperamide, 3mg/kg; <sup>c</sup> compared to 200 mg/kg; <sup>d</sup> compared to 400 mg/kg; <sup>1</sup>p<0.05, <sup>2</sup>p<0.01, <sup>3</sup>p<0.001; 80ME, 80% methanol extract, Negative controls received distilled water.

### In vivo anti-diarrheal index

Results from the determination of *in vivo* ADI revealed that the ADI increased with dose for each fraction and the higher tested dose had the maximum ADI when compared with other tested doses of the extract and the positive control as shown in table 4.

**Table 4:** In vivo anti-diarrheal index of 80ME of leaves of Schinus molle L

| Group | Delay in defecation time (%) | Gut meal travel reduction (%) | No of wet faces reduction (%) | Anti-diarrheal index (ADI) |
| --- | --- | --- | --- | --- |
| 80ME100 | 170.5 | 34.2 | 53.0 | 67.6 |
| 80ME200 | 417.4 | 59.2 | 75.8 | 123.3 |
| 80ME400 | 1303.8 | 73.7 | 87.8 | 203.1 |
| Loperamide | 606.8 | 72.6 | 77.3 | 150.4 |

### Preliminary phytochemical screening

The preliminary phytochemical screening of 80ME revealed the presence of almost all tested constituents except tannins and glycosides as depicted on table 5.

**Table 5:** Preliminary phytochemical screening of the 80% methanol extract of the leaves of Schinus Molle

| **Constitutes** | **80% methanol Extract** |
| --- | --- |
| Cardiac glycosides | - |
| Flavonoids | + |
| Alkaloids | + |
| Saponins | + |
| Steroids | + |
| Tannins | - |
| Terpenoids | + |

## Discussion

Medicinal plants have been used for the treatment of various disorders including diarrhea and related gastrointestinal disorders despite the fact that their safety and efficacy profiles have not been well addressed. It is, therefore, important to properly evaluate the safety and efficacy profile of medicinal plants that are being used in traditional medicines. The need for newer, more effective, cheaper and safer antidiarrheal drugs has become a paramount concern to have safe and cost effective therapeutic alternatives (23).

The present study was aimed to evaluate the antidiarrheal activity of the hydro-alcoholic leave extract of Schinus molle L by using different experimental models of diarrhoea in mice. In all models, diarrhoea was induced by administering castor oil to each mouse. Castor oil produces diarrhoea due to its active metabolite, a ricinoleic acid which is liberated by the action of lipases in the upper part of the small intestine. It facilitates the accumulation of fluid in the intestine and alters the motility of GI smooth muscles (24).

In the castor oil induced diarrhoea model, the extract produced a significant effect on all parameters measured: onset of diarrhoea, the number of wet and total stools and weight of wet stools. This result is in concordance with the report on methanol fraction of the leaves of L. camara (24), the aqueous leaf extract of Leaves barks of L. camara(25), 80% Methanolic Leaf Extract of J. schimperiana (26) and Hydromethanolic Root Extract (27).

The phytochemical analysis of the extract revealed the presence of different bioactive agents. Among the secondary metabolite identified flavonoids and phytosterols are known to modify the production of cyclooxygenase 1 and 2 (COX-1, COX-2) and lipooxygenase (LOX) thereby inhibiting prostaglandin production (26). Tannins present in the extract precipitate the proteins in the intestinal mucosa by forming the protein tannates, which make the intestinal mucosa more resistance to chemical alteration and hence reduce the peristaltic movements and intestinal secretion. Therefore, the anti-diarrheal activity of S. Mole crude extract observed in this study may be attributed to the presence of flavonoids, alkaloids, tannins and phytosterols in the crude extract.

To settle the antidiarrheal activity of leaves of S. molle the possible mechanism of action was tested on intestinal motility and enteropooling models.

The enteropooling model was designed to assess the antisecretory effect of hydromethanolic leaf extract of S.Molle In this model; the extract significantly reduced the intraluminal fluid accumulation when compared to the negative control. This result was in line with the results of other studies (24–27). The active metabolite of castor oil, ricinoleic acid, induces irritation and inflammation of the intestinal mucosa, leading to release of prostaglandins. The prostaglandins thus released stimulate secretion by preventing the reabsorption of sodium chloride and water. Thus, it is possible that the extract significantly inhibits gastrointestinal hypersecretion and enteropooling by increasing reabsorption of electrolytes and water or by inhibiting induced intestinal accumulation of fluid (27). The anti-enteropooling activity of the extract could also probably be related to the existence of phytochemical constituent including flavonoids, steroids and tannins (27–29).

In the castor oil-induced gastrointestinal motility model, it was observed that the extract significantly suppresses the movement of the charcoal marker at all tested doses of the extract (100 mg/kg, 200 mg/kg, and 400 mg/kg) as compared to the negative control. The higher percentage of inhibition (66.2%, p < 0.001), of the marker perceived at maximum dose was almost comparable to loperamide (64.5%, p < 0.001 at the dose of 3 mg/kg). This finding showed that the extract can influence the peristaltic movement of the intestine thereby indicating the presence of intestinal antimotility activity. Concerning this, several plants have shown antidiarrheal activities by reducing the gastrointestinal motility and its secretions (24–29).

Several studies suggested that the anti-motility properties of herbs are mostly due to flavonoids; by inhibiting the release of autacoids and prostaglandins results in inhibiting motility and hydro-electrolytic secretions induced by ricinoleic acid. Tannins may also show an anti-motility effect by reducing intracellular Ca2+ through decreasing Ca2+ inward current or increasing calcium outflow and finally resulting in reducing peristaltic movement and intestinal secretions due to induction of the muscle relaxation (28). Pre-treatment with 80% methanol extract significantly reduced peristaltic movements as evidenced by the decrease in the distance travelled by a charcoal meal in the GIT, showing that these crude extracts could have anti-motility activity due to flavonoid and tannin constituents

Like the castor oil induced and enteropooling diarrheal model, maximum effect was observed with the highest dose of the extract rather than the standard drug in charcoal meal test. This might be due to different secondary metabolites in the extract that may prolong the time for absorption of water and electrolytes through hampering the peristaltic movement of the intestine.

Clinically, diarrhoea may result from disturbed bowel function, in which case, there is impaired intestinal absorption, excessive intestinal secretion of water and electrolytes and a rapid bowel transit (25). In vivo, ADI is a measure of the combined effects of the different components of diarrhoea, including purging frequency, the onset of diarrheal stools and frequency of intestinal movement. Besides, higher ADI value is a measure of the effectiveness of an extract in curing diarrhoea. The ADI value increased with dose, suggesting the dose dependency of this parameter. The highest selected dose of the extract, with the highest ADI value, is endowed with the best antidiarrheal activity when compared with other selected doses as indicated on the above results.

## Conclusion

The results of this study revealed that the hydro-alcoholic leave extract of schinus molle endowed with significant antidiarrheal activity. It inhibited the frequency of defecation and reduced greatly the wetness of faecal excretion.

Moreover, it also produces an inhibitory effect on castor oil induced intestinal secretion and gastrointestinal propulsion. These antidiarrheal activities of the extract may be attributed to the presence of phytochemicals including tannins, alkaloids, saponins, flavonoids and phytosterols that act individually or collectively. These findings provide a scientific support for a traditional use of the leaves of Schinus molle as diarrhoea remedy.

## Acknowledgment

The financial support of Wollo University is gratefully acknowledged.

